# Impact of secondary pests on carbon allocation in declining beech trees after the 2018-2020 drought episode

**DOI:** 10.64898/2026.07.22.739995

**Authors:** Gaertner Pierre-Antoine, Bréda Nathalie, Gérard Bastien, Levillain Joseph, Schmuck Hubert, Larousse Tony, Badeau Vincent, Saintonge François-Xavier, Massonnet Catherine

## Abstract

1. During the extreme drought period of 2018-2020, secondary pest attacks were observed in some declining beech trees (*Fagus sylvatica*). The main species identified are *Agrilus viridis* and *Taphrorychus bicolor*, which are known to attack beech trees more frequently after periods of intense drought. The aim of this study is to investigate if and how these two cambium-feeding insects in interaction with drought contribute to beech decline. We assessed the modification of the carbon reserves stock and carbon allocation to growth and fruiting in declining beech trees.
2. Past radial growth, non-structural carbohydrates (NSC) concentrations in several organs and the quantity of beechnuts were measured in 38 beech trees with contrasting levels of biotic attack in 2020. Total amount of carbon reserves was calculated on the basis of NSC concentrations, and the amount of carbon allocated to growth and fruiting at the tree level was assessed using allometric relationships.
3. Beech trees most severely attacked by secondary pests showed lower radial growth for last ten years than trees that have not been attacked. NSC concentrations and the amount of carbon reserve in the coarse roots and stems of trees were also lower in beech trees with intense biotic symptoms compared to unaffected trees. However, the amount of carbon allocated to growth and fruiting during the 2020 growing season did not differ significantly among biotic attack classes.
4. *Synthesis:* Regardless of biotic attack intensity, declining beech trees exhibited abnormally low carbon reserves. Trees most severely affected by insects in 2020 appeared to be the most vulnerable, as evidenced by reduced radial growth over the past decade. Future experiments are needed to explore further whether insects impacted directly carbon reserves, or whether low carbon reserves in trees is a vulnerability factor promoting insects attacks. Finally, declining beech trees allocated a similar amount of carbon to fruiting as reported in healthy stands, suggesting that fruiting is a prioritised carbon sink, even under severe drought and biotic attack.

## 1 Introduction

From 2018 to 2020, central European forests were exposed to three consecutive extreme drought periods, which induced the decline of many tree species. One of the most affected species was the European beech (*Fagus sylvatica* L.). From 2019 onwards, heavy leaf loss, branch mortality and dead trees were observed in many places (Schuldt et al. 2020; Obladen et al. 2021; Frei et al. 2022). In some cases, declining trees were also attacked by buprestids and bark beetles (Frei et al. 2022; Mirabel and Gaertner 2023), mainly *Taphrorychus bicolor* and *Agrilus viridis*. These insects are considered as opportunistic secondary pests, which means they preferentially attack weakened trees, causing significant damage and accelerating their decline (Asthalter and Lehmann 1979; Lakatos and Molnár 2009; Brück-Dyckhoff et al. 2017; Brück-Dyckhoff et al. 2019). Both extreme droughts and biotic attacks can induce strong physiological impairments, including changes in carbon metabolism (Anderegg et al. 2015; Huang et al. 2020).

Under non-stressful conditions, many vital functions mobilise carbon. A major part of the assimilates produced during photosynthesis is consumed in living cells via mitochondrial respiration (Granier et al. 2008; Hartmann and Trumbore 2016). This carbon is also invested for growth (Smith and Stitt 2007; Klein and Hoch 2015) and reproduction (Hoch and Keel 2006; Hoch et al. 2013). Assimilated carbon is used to produce more complex molecules such as phenols and alkaloids, which are used in particular for defence and communication (Kesselmeier et al. 2002; Huang et al. 2020). Finally, some of the carbon is stored, which allows to buffer the gap between carbon availability and the tree’s needs (Chapin et al. 1990; Sala et al. 2011). Carbon is mainly stored in the form of non-structural carbohydrates (NSC). In beech trees, carbon reserves are mainly composed of starch, glucose, fructose and sucrose (Hoch et al. 2003). These reserves are most concentrated in the branches and roots, but the stem is the main storage organ due to its large biomass of living tissue (Barbaroux et al. 2003). In beech trees, these reserves are mobilised in spring to produce leaves, flowers and primary growth (Barbaroux and Bréda 2002; Hoch et al. 2013). After leaf expansion, the main carbon sinks are secondary growth and respiration, which are mainly fuelled by recent photosynthetic products (Epron et al. 2012). During the growing season, the amount of assimilates is also co-allocated to storage to restore carbon reserves, which are at their highest in autumn (Barbaroux and Bréda 2002; Epron et al. 2012). In contrast, fruit production does not depend on reserves but on recent photosynthates (Hoch and Keel 2006; Hoch et al. 2013; Ichie et al. 2013). Beech is a masting species, meaning a synchronised occurrence among trees of large amounts of fruits and seeds at irregular intervals. Some years, fruiting can therefore be a major carbon sink, particularly in mature beech trees (Genet et al. 2010). This sink then competes with growth for carbon allocation, but in non-stressful situations, fruiting causes only a slight or no decline in radial growth (Mund et al. 2010; Hacket-Pain et al. 2017).

Moreover, stressful events such as soil water deficits, caterpillar defoliations or late frosts induce changes in carbon allocation. For example, when the relative water reserve in the soil falls below a threshold of 40%, trees, including beech trees (Lemoine, Cochard, et al. 2002; Granier et al. 2007), reduce their transpiration by closing their stomata. Hydraulic conductance decreases due to the limitation of leaf water potential (Bréda et al. 1993), and embolism formation in the xylem vessels can sometimes occur and further increase hydraulic resistance (Sperry et al. 1998). If water stress persists, the embolism may spread to other vessels, which can cause the death of the affected organ or even the tree, as observed in beech saplings (Barigah et al. 2013). In forest beech trees nevertheless, a segmentation of hydraulic properties depending on light exposure of branches (Lemoine, Cochard, et al. 2002) with a high plasticity in case of light exposure change (Lemoine, Jacquemin, et al. 2002), is an efficient property to avoid propagation of embolism. The stomatal closure also reduces carbon assimilation (Bréda et al. 2006), and therefore the amount of assimilates available to the tree. Carbon is then allocated primarily to metabolism (Hartmann and Trumbore 2016) and storage, to the detriment of growth (Chuste et al. 2020; Huang et al. 2021). Radial growth in beech trees is known to be significantly reduced when rainfall is low and temperatures are high between May and July, with a lag effect the following year (Badeau 1995; Lebourgeois et al. 2005; Scharnweber et al. 2011). On the contrary, NSC reserves are generally maintained at normal levels during droughts (Martínez-Vilalta et al. 2016; Dietrich et al. 2019; Piper and Paula 2020) or even temporarily increased (Chuste et al. 2020). Beechnut production also appears to be maintained at normal levels during water deficits (Hesse et al. 2021), at the expense of growth, as a more pronounced decline in radial growth is observed when masting coincides with drought (Hacket-Pain et al. 2017). However, during extreme and/or repeated water deficits, especially inducing premature leaf fall as in 2003 or 2018, the amount of carbon assimilated becomes so limited that storage can no longer be maintained at a normal level (Bréda et al. 2006). Carbon reserves may thus decrease (Hartmann et al. 2013; Trifilo et al. 2017) or even collapse, potentially causing tree death (Galiano et al. 2011; Chuste et al. 2020). An abnormally low production of beechnuts was also observed during the extreme drought and heat summer of 2018 (Nussbaumer et al. 2020).

The malfunctions caused by drought then promote successful attacks by pests, which overcome the host tree’s induced defence mechanisms and manage to colonise it (Gaylord et al. 2013; Netherer et al. 2015; Kolb et al. 2016), eventually leading to its death (McDowell et al. 2011). For example in conifers, insects that feed on cambium can themselves cause a decrease in carbon reserves, particularly in drought conditions (Wiley et al. 2016; Erbilgin et al. 2021). Water deficits and biotic attacks would thus induce trade-offs in the carbon allocation to growth, reproduction, storage and defence functions.

Biotic and abiotic hazards are often associated in a cascade of events in forest decline (Sinclair and Hudler 1988; Manion 1991; Landmann 1994; McDowell et al. 2011). Abiotic hazards generally act as inciting factors, whose impacts on tree physiology (growth, carbon reserves and potentially reduced defence mechanisms) increase their vulnerability to biotic attacks, acting as contributing factors to tree death. However, the physiological mechanisms involved in this vulnerability of trees to biotic attacks remain poorly understood. This prior weakening of trees is often necessary for insects to colonise them (Anderegg et al. 2015). Although the mechanisms by which cambium-feeding insects select their hosts are still poorly understood, they likely prioritise the most suitable trees to maximise their fitness (Raffa et al. 2016; Lehmanski et al. 2023). Hence, in silver fir and white fir trees, radial growth is lower in trees attacked by bark beetles, regardless of their size or status (Durand-Gillmann et al. 2014; Stephenson et al. 2019). Outside of drought conditions, it has also been reported that oak trees with lower winter starch reserves were more heavily attacked by bark beetles the following spring (Dunn et al. 1987; Dunn and Potter 1990).

Most studies examining the combined effects of drought and insect attacks on tree physiology focus on conifers. Much remains to be discovered about angiosperms, particularly their carbon allocation strategy in response to cumulative water and biotic stress. In this study, we investigated the impact of secondary pests interacting with extreme water deficits on carbon management in mature beech trees. Radial growth and NSC concentrations in different organs were measured on beech trees with contrasting crown condition and density of biotic attack. As suggested by Wiley (2020), the survival of trees under stress may be more related to the amount of carbon available in the organism than to the concentration measured in its tissues. We also estimated carbon quantity in order to better understand its use by trees and its allocation to different sinks. The intensity of the water deficit experienced by trees on the plot was also quantified using the Biljou© water balance model (Granier et al. 1999). Our hypotheses are: I) past radial growth of beech trees heavily attacked by insects is lower than that of asymptomatic trees, confirming the preferential attack of weakened individuals, II) NSC concentrations are lower in trees attacked by insects, III) trees undergoing biotic attacks reduce their reserve carbon stocks and carbon allocation to growth, but maintain allocation to fruiting.

## 2 Materials and Methods

### 2.1 Stand description

This study was conducted on a mature beech (*Fagus sylvatica* L.) stand in Bliesbruck communal forest in north-eastern France (49°6’32’’ N, 7°9’31’’ E, mean elevation 315 m). Mean annual temperature and precipitations during the 1959-2017 period were 9.5 °C and 901 mm respectively. Six plots were uniformly installed across the stand to measure dendrometric variables (basal area, tree density, mean diameter) and the leaf area index (LAI) at the centre of each plot in the four cardinal directions using a LAI-2000 (Li-Cor, Lincoln, Nebraska, USA) fitted with a three-quarter closed mask. This stand was an even-aged high forest almost pure in beech, whose mean tree height was 34 m and stand density was 98 trees ha^-1^. Soil characteristics were slightly different across plots, consisting in hypereutric cambisol and epistagnic luvisol. Despite this difference, extractable soil water was homogenous across the stand, ranging from 153 to 163 mm on about 100 cm depth. From 2019 onwards, a decline in beech trees was observed, characterised by secondary pest attacks and by symptoms of crown degradation i.e., high leaf loss and branch mortality.

### 2.2 Description of trees and their level of insect attack

In September 2020 (8-10 September), 38 mature beech trees presenting contrasting crown conditions were selected across the stand among the trees harvested by the French National Service of Forests (ONF). Consequently, sample collection was not allowed on every tree to not impair wood quality for commercialisation, which explains that the number of sampled trees varies among the analyses (see 2.3 and 2.4). Mean age of the sampled trees was 156 ± 11 years as measured by tree coring. The circumference and the height of each tree was measured. Before harvest, the crown transparency of the trees was evaluated on standing trees according to the ICP Forests leaf loss assessment protocol (Eichhorn et al. 2016). Once they have been felled, the presence of insect attacks (galleries, emergence holes, bleeding cankers, sawdust) was detected first on the bark and then, after gentle stripping, from the base of the stem to the top of the tree, including large branches. Most of the trees with biotic symptoms showed both active galleries and emergence holes (**FIGURE S1**) resulting from infestations in previous years (probably from 2019), so current and past attacks (less than two years) were not distinguished in the analyses. The number of emergence holes and galleries, the presence of seepages and the stem length concerned by these symptoms were assessed. The species responsible from these symptoms were determined. Most of the trees with biotic symptoms were attacked by both *Agrilus viridis* and *Taphrorychus bicolor*, so, species were not distinguished in further analysis. This information was aggregated into a 3-classes notation describing the extent and the intensity of the biotic symptoms: A = not significantly attacked (no or few isolated emergence holes); B = moderately attacked (scattered emergence holes and galleries covering a stem length longer than one meter); C = heavily attacked (numerous emergence holes and galleries covering the stem over several meters).

### 2.3 Dendrochronological analysis

A core was sampled on 36 trees with a 5-mm core drilling machine, for dendrochronological analyses. 16 trees were sampled at 1.3 m, with the remainder sampled at 11 m (logs for which sampling was not permitted). Eight trees were sampled at both 1.3 m and 11 m in order to establish a relationship between radial growth at these two heights. After surface treatment with a microtome (Gärtner and Nievergelt 2010), tree-ring widths were measured with an incremental encoder (Velmex Inc., Bloomfield, NY, USA) coupled with the MeasureJ2X program. Then, trees were visually cross-dated using CDdendro software (Cybis Elektronik & Data AB, Saltsjöbaden, Sweden). Ring width were converted to basal area increment (BAI) using the ‘dplR’ R package (Bunn 2008).

According to the Pressler’s law (1864), cited by Larson (1963), ring surface is constant along the stem up to the first living branch. However, this law was not verified in trees sampled at both 1.3 m and 11 m, even if this difference was not significant any more from 2000 onwards. To be able to compare growth trends between trees sampled at 1.3 m and at 11 m, growth was detrended according to tree age and the height of sampling using a relationship between BAI and cambial age (see Supplemental S3). First, the BAI was averaged for each cambial age available in the data set according to the height of sampling. Then, a LOESS regression (span = 0.8) was fitted to the relation BAI = f(cambial age) for each sampling height. The LOESS regression was set to a fixed value for cambial ages greater than 125 years at 1.3 m and greater than 100 years at 11 m as the BAI curve becomes unstable at high cambial ages. Raw BAI was finally divided by the value of the predicted BAI according to the cambial age of the tree ring and to the height of sampling to calculate a growth index:

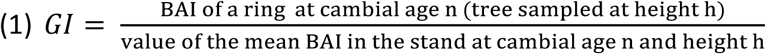

### 2.4 Sampling for carbohydrates measurements

Thirty-three trees were sampled after the cutting in early autumn (21-25 September 2020) because this period corresponds to the maximum of NSC reserve concentrations and amounts (Barbaroux and Bréda 2002). The samples were collected from all tree compartments (coarse roots, stem and branches including basal section, twigs). The samples from the stem were taken with a 5-mm increment borer at three different heights (0.3, 11 and 22 m). Samples were also collected at the base of three branches, whose mean diameter was 2.4 ± 0.7 cm. For each branch, two twigs from 2020 and 2019 growing seasons and all the nuts were harvested. Furthermore, samples were collected from three coarse roots about 40 cm from the stem (**FIGURE 1**). The crown of four severely declining trees exploded during their felling, so it has been impossible to properly sample these trees in the stem at 22 m high and to harvest branches.

**FIGURE 1.**
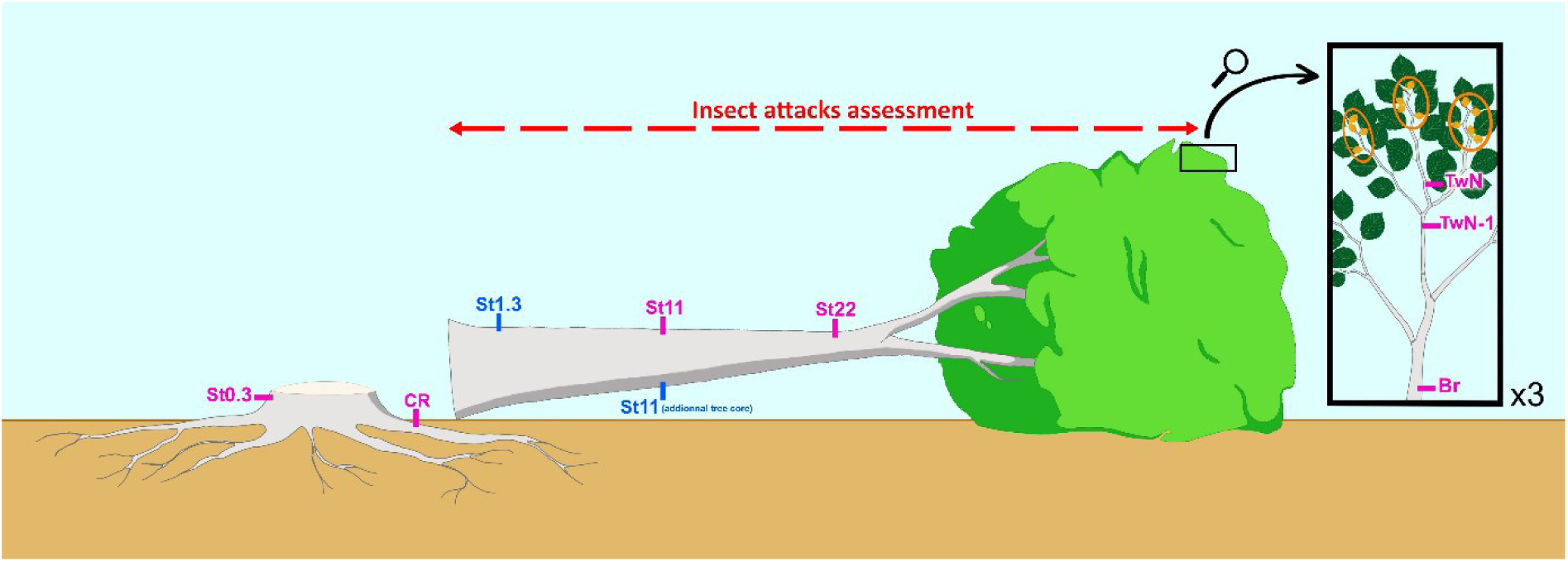
Sampling scheme for carbohydrate measurements and radial growth analysis. CR: coarse roots; St: stem, followed by the coring height (0.3, 1.3, 11 and 22 m); Br: branches; TwN-1: twigs from the 2019 growing season; TwN: twigs from the 2020 growing season. Samples from coarse roots, branches and twigs collected in triplicate. Samples dedicated to carbohydrate measurements are noted in pink and those for radial growth analysis in blue. The box presents, in orange, beechnuts and fruit husks collected in branches to quantify fruiting.

Immediately after sampling, tree cores were wrapped in plastic protection and conserved in a cooler. The following day in the laboratory, they were cut in 2-cm sections, weighted, frozen and stored at - 80 °C, until freeze-dried. All the samples were freeze-dried (FreeZone® 12L −84 °C with Console Freeze Dryers with Stoppering Tray Dryers, Labconco, USA) and finely ground in a ball mill (Mixer Mill MM400, Retsch, Germany) for tree core sections and in a cup mill (CEPI SODEMI CB2200, Cergy, France) for other samples. The samples were stored in airtight containers until carbohydrate analysis.

We distinguished the inner and outer parts of the stem cores based on soluble sugar content. We defined the inner part as beginning below the concentration threshold of 1% soluble sugars (per gram of dry matter). Based on preliminary analyses of 2-cm sections, this outer-inner boundary was set at 10 cm (the first five sections after the bark) for the stump core and at 6 cm (the first three sections) for the two other stem cores. 2-cm sections from the inner part were ground and assayed together, outer sections were assayed individually.

The samples from branches and nuts were weighted, and an aliquot was collected from these samples and weighted. These aliquots were dried in an oven at 80°C for three days, then weighted to determine dry weight. The other samples were dried, ground and stored in airtight containers.

### 2.5 Carbohydrate analysis

Soluble sugars were extracted from 10 to 15 mg powder from the ground core samples mixed into 0.5 mL of 80 % ethanol (v/v) and incubated for 20 min at 80 °C. Extraction was repeated twice and the three supernatants were collected in a tube and evaporated to dryness under vacuum (SpeedVac: Refrigerated CentriVap Vacuum Concentrators, Labconco). The resulting dry residue was resuspended in 1.5 mL ultrapure water by sonication and agitation and then stored at −20°C. Glucose, fructose and sucrose concentrations were successively measured following the protocol described in Gomez et al. (2007), in which the assay is based on the conversion of glucose, fructose and sucrose into glucose-6-phosphate by enzymatic hydrolysis, proportional to the reduction of NAD to NADH. Assays were carried out in 96-well microplates in triplicate using an automated system (Microlab Starlet pipetting station, Hamilton, Switzerland) for sample collection and reagent dispensing. A standard series of known concentrations of these three sugars was prepared on each microplate. Blanks (samples containing the reaction medium) without the addition of enzymes were also prepared. After adding a reaction buffer containing NAD and ATP, the enzymatic reagents (1) hexokinase (Roche 11426362001) and glucose-6-phosphate dehydrogenase (Roche 10165875001), (2) phosphoglucose isomerase (Sigma P5381) and (3) β-fructosidase invertase (Sigma I9274) were added sequentially, followed by 3-hours incubations at room temperature in the dark. The subsequent increase in NADH was measured at the specific wavelength of 340 nm with a microplate reader spectrophotometer (Xénius XMA – SAFAS, Monaco) and used to determine sugars concentrations.The glucose, fructose and sucrose concentrations were summed as soluble sugars.

The pellet remaining after the soluble sugar extraction was dried under vacuum. The starch was extracted from the dried pellets in 1.5 mL of a 0.2 M KOH solution incubated for 20 min at 80 °C. Starch contents were determined after hydrolysis into glucose by amyloglucosidase (Sigma A1602) for 2h at 50° in sodium acetate buffer. Briefly, the obtained glucose was determined colorimetrically in 96-well microplate format using an enzymatic reagent containing glucose oxidase (Sigma G6125) / peroxidase (Sigma P8125) and orthodianisidine (Sigma D9143). After 10 min incubation in the dark and adding 6N hydrochloric acid, absorbance was measured at 530 nm with a microplate reading spectrophotometer (Xénius XMA – SAFAS, Monaco), with glucose as a standard (Chow and Landhausser 2004). Starch was expressed as equivalent glucose. Starch, soluble sugars and their sum (total NSC concentrations) were expressed as grams per 100 g of dry sample mass (g 100 g^-1^_DM_).

### 2.6 Estimation of the quantity of carbon stored or allocated to fruiting and growth at tree level

To estimate the quantity of carbon stored, the biomass of each organ was multiplied by its NSC concentration expressed in glucose equivalent and by 0.4, which corresponds to the carbon content of glucose. The biomass was estimated using equations (the stem was considered as a truncated cone) and allometric relationships (from Genet et al. (2011)). The complete method is detailed for each organ in the supplemental section S2.

To estimate the quantity of carbon allocated to aboveground radial growth, the volume of the tree-ring produced in 2020 was multiplied by the density (566 g dm^-3^) and the carbon content (46 %) of beech wood (Barbaroux et al. 2003). The detailed procedure is given in the supplemental section S2. To estimate the quantity of carbon allocated to fruiting, the dry mass of beechnuts and beechnut husks (the calculation is detailed in S2) was multiplied by their carbon content (50 %) (Lebret et al. 2001).

Unlike growth and fruiting, carbon allocation to storage in 2020 could not be measured because this required measuring NSC concentrations in each compartment twice, in spring and in autumn, to measure the difference in NSC reserves between their lowest and highest level (Barbaroux and Bréda 2002), which was impossible with our sampling method.

### 2.7 Water Deficit Index calculations

Soil water deficit was quantified with the [BILJOU©](https://appgeodb.nancy.inrae.fr/biljou/en/) forest water balance model (Granier et al. 1999) which calculates the daily dynamics of elementary water fluxes (transpiration, rainfall interception, runoff) and the resulting soil water reserve. The calculations were performed retrospectively to compute the intensity of the past drought events experienced by the stand. The Water Deficit Index (WDI) was calculated as the sum of the daily differences between the soil relative extractable water (REW) and 0.4, divided by 0.4. A REW value of 0.4 is considered to be the threshold for soil water deficit (Black 1979; Granier et al. 1999), and is also verified for beech stands (Granier et al. 2000). WDI is dimensionless and can be considered on a monthly, seasonal or annual basis or summed over several years.

The model requires stand characteristics (budburst and leaf fall dates, maximum leaf area index) and soil properties (depth of soil rooted layers and for each soil layer: proportion of fine roots, extractable water, soil bulk density, soil water content at the permanent wilting point). Daily meteorological data are also required (total rainfall, mean wind speed, mean temperature, mean relative air humidity and global radiation to compute potential evapotranspiration).

Bud burst date was set at day of year 112 and leaf fall date at day of year 300, which corresponds to averaged values for beech in north-eastern France (Lebourgeois et al. 2002).

As the extractable soil water only differs by 10 mm between plots, which has no significant influence on the soil water balance, only one parametrisation for the whole stand was chosen. The soil characteristics of the cambisol were used as this soil type was predominant in the stand. A LAI of 5.8 m^2^ m^-2^ was chosen, corresponding to the average LAI in the stand measured in 2020.

### 2.8 Statistical analysis

All the analyses were performed in R (R Core Team 2022). To study the annual radial growth differences among biotic attacks classes, GI was first log-transformed as the standardised residuals of GI did not verify the normality and the homoscedasticity. Then, linear mixed-effect models (‘lme’ function of the ‘nlme’ R package) were fitted for GI with the year, the biotic attack class and their interaction, and with individual as a random effect. This procedure was followed by a post-hoc test with the ‘glht’ function. To test differences among biotic attacks classes for the other variables examined (carbohydrate concentrations, carbon quantity, dendrometric variables and averaged radial growth), the generalised least square method was used with the ‘gls’ function of ‘nlme’ R package (Pinheiro et al. 2023). This procedure was followed by a post-hoc test with the ‘glht’ function from the ‘multcomp’ R package (Hothorn et al. 2008). This procedure was chosen because it allows us to take into consideration the structure of the intra-group heteroscedasticity and is more effective to compare groups with small numbers.

## 3 Results

### 3.1 Dendrometry, leaf loss and radial growth of trees according to the intensity of biotic attacks following the severe drought 2018-2020

No difference in diameter or height was observed among biotic attack classes (FIGURE S4), so the dendrometric characteristics of the trees do not bias the constitution of the biotic attack classes. On the other hand, trees attacked by insects have a higher average leaf loss than trees without biotic symptoms, but the difference is only significant for moderately attacked trees (FIGURE S4).

No significant difference in annual radial growth was detected among biotic attack classes (**FIGURE *2*a**). Regardless of biotic symptoms class, radial growth has followed a downward trend since the 1990s and with a GI systematically lower than 1 since 2014. This decline started earlier and appears to be more pronounced for class C since 2008, but this difference among classes does not appear to be significant. The chronologies show several past growth crises linked to sequences of consecutive soil water deficits (sequences 1989-1990-1991 and 2003-2005) (**FIGURE 2b**). Radial growth declined significantly compared to the previous five years during the periods 1989-1992 (−34%) and 2003-2005-2006 (−27%). The soil water deficit cumulated during the three consecutives growing seasons 2018-2019-2020 are the more intense of the 60’s last years.

**FIGURE 2.**
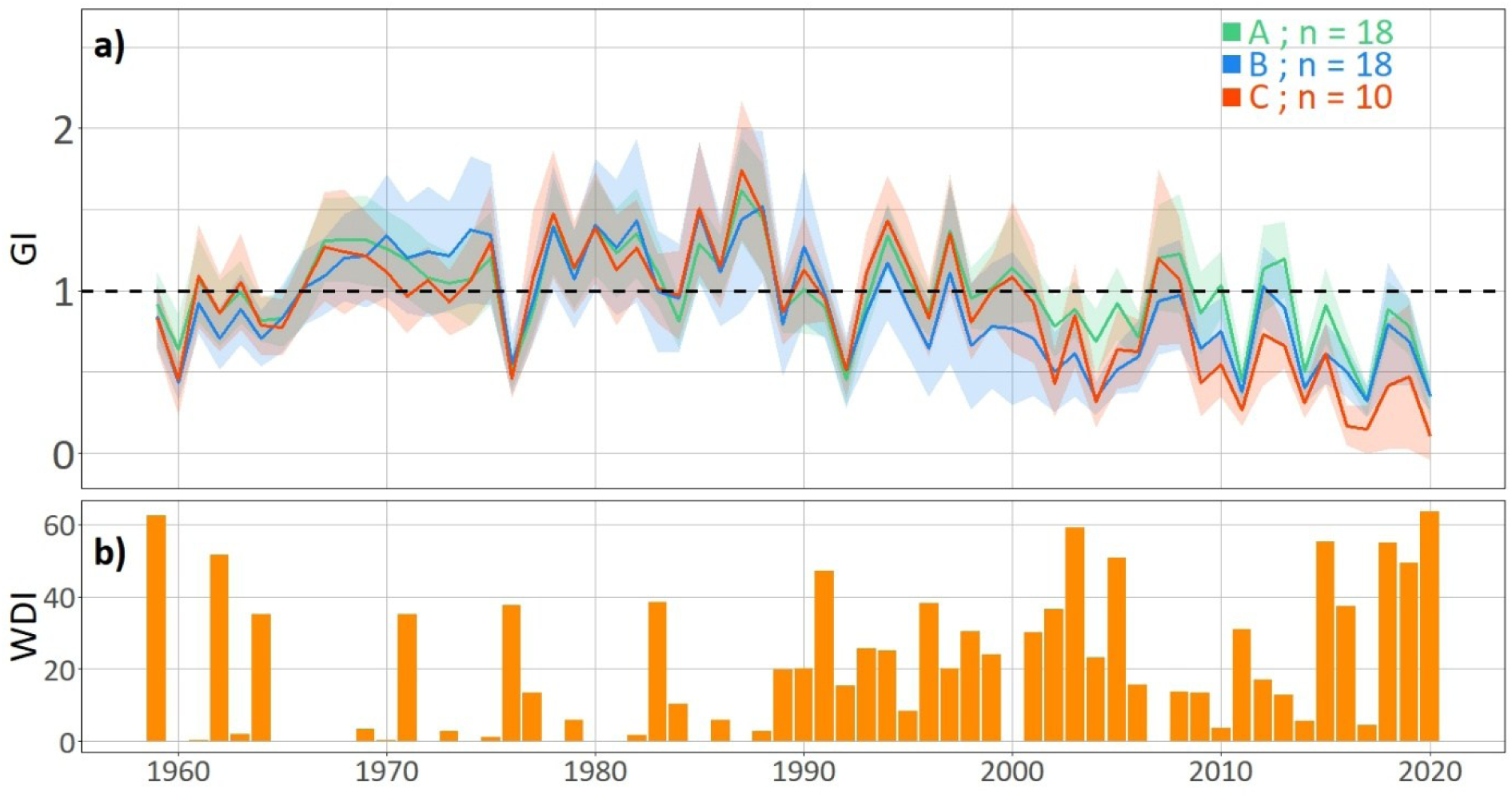
**a)** Mean yearly growth index (GI) from 1959 to 2020 according to the biotic attacks class (A = not significantly attacked; B = moderately attacked; C = heavily attacked). The shaded areas around the lines represent the standard error and the dashed horizontal line to 1 indicate the mean growth of all trees in this forest detrended of the age effect. **b)** Yearly water deficit index (WDI) for the study site.

To compare average growth over time, **FIGURE 3** distinguishes three-year periods starting from 2006-2008 to the recent period of 2018 to 2020, which saw a recurrence of severe water deficits. This allows detecting a significant radial growth decline in the most attacked trees in comparison to the non-attacked trees, significant decline beginning in the period 2009–2011. This is consistent with the radial growth chronologies presented in **FIGURE 2a**, even though the tests carried out for this analysis did not detect year-by-year significant differences. From 2009 to 2011, the radial growth of trees moderately attacked by insects in 2020 was intermediate between that of undamaged trees and that of heavily attacked trees (**FIGURE 3**).

**FIGURE 3.**
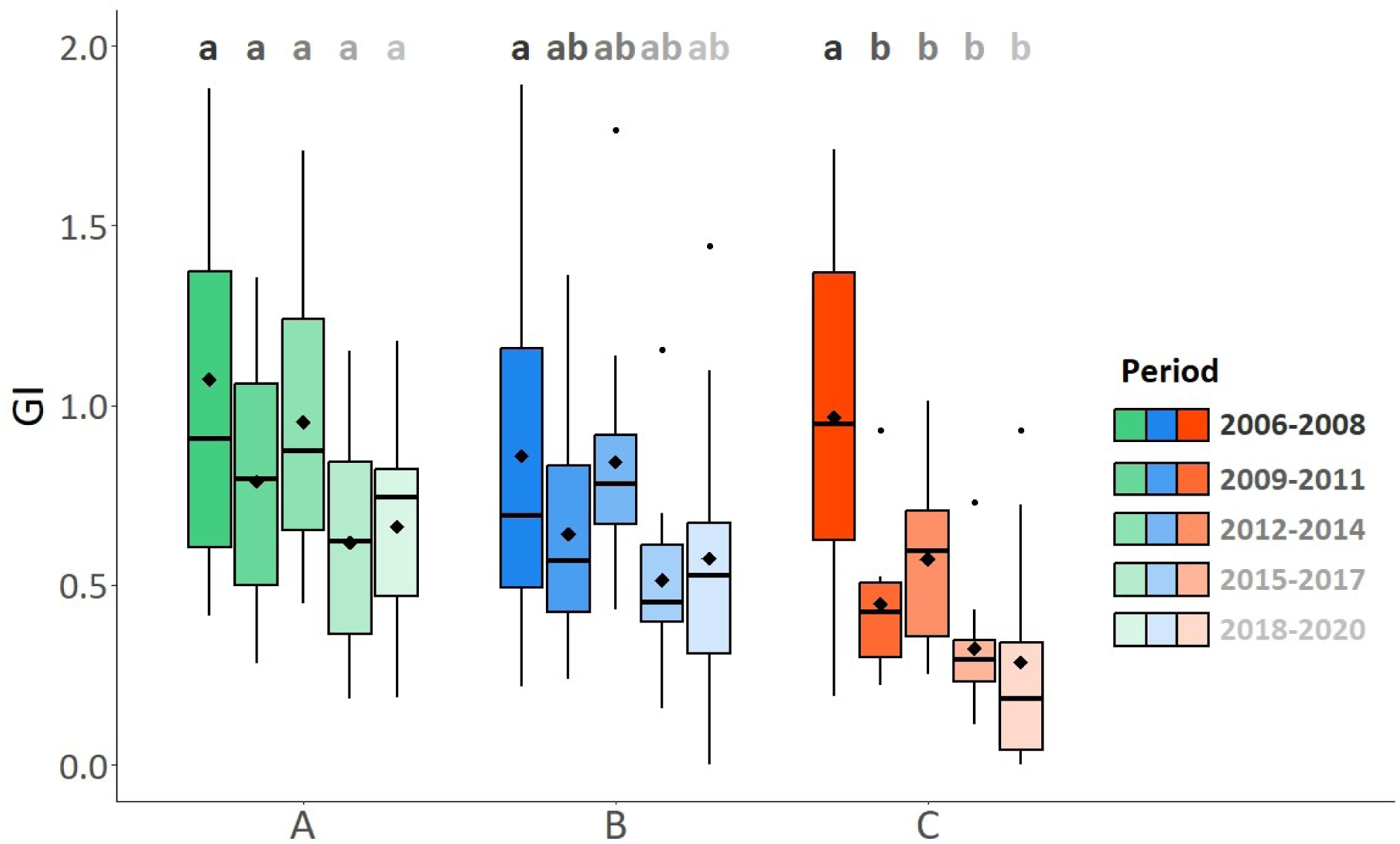
Average 3-year growth index (GI) for five periods ranging from 2006-2008 to 2018-2020 according to the biotic attacks class (A = not significantly attacked; B = moderately attacked; C = heavily attacked). In boxes, diamonds represent the mean, lines represent the median. Different letters represent a significant difference (p < 0.05) in GI among biotic attacks classes for each year or period.

### 3.2 Differences in NSC concentrations among biotic attacks classes

NSC concentrations strongly varied from one compartment to another (**FIGURE 4**). The branches and particularly the recent twigs showed the highest NSC concentrations (7.8 and 7.4 g.100 g^-1^ on average for those in 2020 and 2019 respectively). NSC concentrations were the lowest in the trunk, particularly at 11 m and 22 m high (1.0 g 100 g^-1^ on average from 0 to 2 cm depth) and decreased radially (at 0.3 m, concentrations fell from an average of 2.3 g 100 g^-1^ from 0 to 2 cm under the bark, to 0.3 g 100 g^-1^ from 10 cm depth to the pith). The concentrations of the different carbohydrates also differed from one compartment to another, with sucrose being the predominant carbohydrate in the twigs but starch dominating in the branches and the outermost section of the trunk at 0.3 m (starch/total soluble sugar ratios of 0.9 and 0.8 respectively). Sucrose and starch concentrations were lower in the coarse roots than in the aerial organs.

**FIGURE 4.**
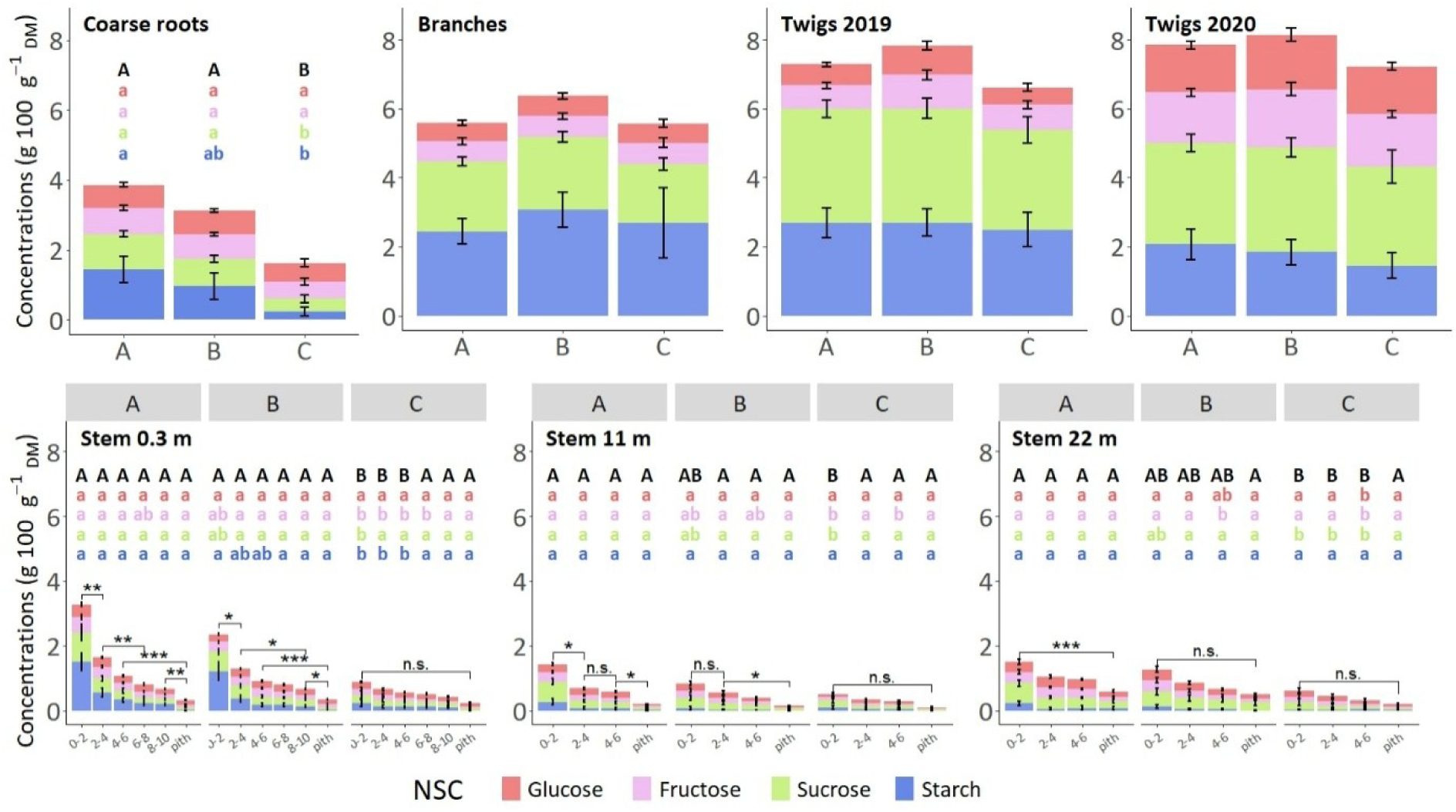
Glucose, fructose, sucrose and starch concentrations in various tree organs according to the biotic attacks class (A = not significantly attacked; B = moderately attacked; C = heavily attacked). For stem, the radial variation of NSC concentrations has been quantified by 2-cm sections from the bark up to 10 cm deep at 0.3 m high, and up to 6 cm deep at 11 m and 22 m high. Beyond that, the inner part of the core is considered as one section: pith. The error bars represent the standard error. Each biotic class A, B, C presented sampling number N = 12, 13 and 8, respectively, for coarse roots, stem at 0.3 m and 11 m. N = 11, 12, and 6 for other compartments because tree crown exploded during their felling, so they were impossible to sample above 11 m high. Different letters represent a significant difference (p < 0.05) in NSC concentrations among biotic attacks classes. The colour of lowercase letters corresponds to that of the different NSCs (glucose, fructose, sucrose and starch), black capital letters correspond to total NSC concentrations (glucose + fructose + sucrose + starch). For stem, asterisks represent significant difference (p > 0.05: n.s.; p < 0.05: *; p < 0.01: **; p < 0.001: ***) in NSC concentrations among radial sections. To improve readability, all relationships between sections were not represented. As NSC concentrations decrease from bark to pith, if, for example, there is a significant difference between sections 6-8 and pith and also between sections 8-10 and pith, only the latter will be represented.

The impact of biotic attacks on NSC concentrations differed among organs. Branches and twigs showed no difference in NSC concentrations according to the intensity of biotic attacks. On the other hand, the trees that were most attacked by insects (class C) had significantly lower NSC concentrations in the coarse roots and the stem, whatever the height of coring, and in particular at the base of the stem (**FIGURE 4**). In the stem, the differences were most apparent in the first few centimetres below the bark, whatever the degree of biotic attack. The differences in NSC concentrations observed were mainly due to lower starch and sucrose concentrations in the attacked trees, whereas fructose and especially glucose concentrations varied little from one class to another or radially. In the coarse roots and the outermost part of the base of the stem, the ratio of starch to soluble sugars fell from 0.6 and 1 respectively in trees that were not attacked by insects to 0.1 and 0.3 in trees heavily attacked by insects.

### 3.3 Carbon stored and allocation to growth and fruiting at the tree level

Total biomass as well as root, stem and branch biomass showed no significant differences among biotic attacks classes (**FIGURE 5**). The stem is by far the organ with the highest biomass (around 55% of total tree biomass), followed by the coarse roots (around 35%) and then the branches (less than 10%).

**FIGURE 5.**
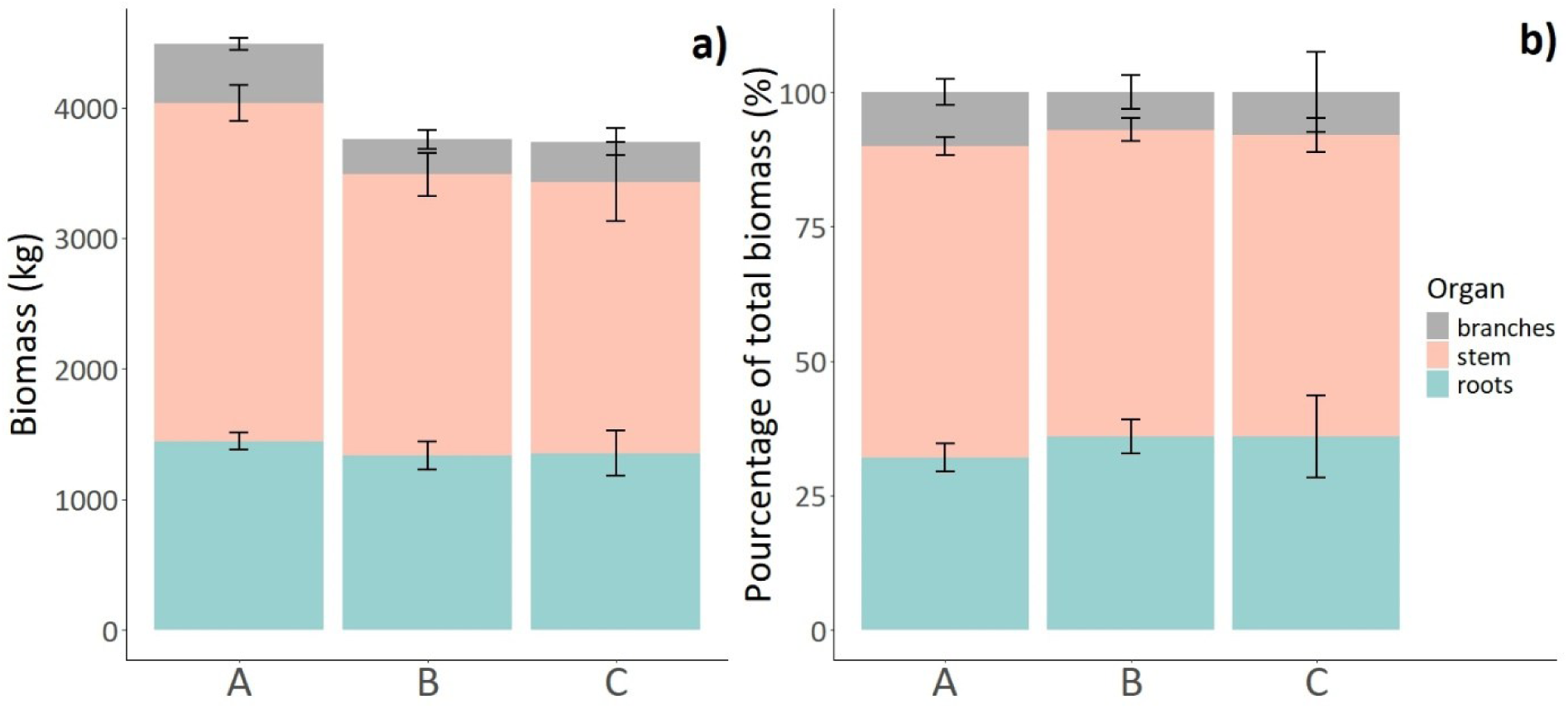
Biomass at the tree level **a)** in kg by organ (branches, stem, roots), **b)** in proportion of total tree biomass according to the biotic attacks class (A = not significantly attacked; B = moderately attacked; C = heavily attacked). No significant difference has been observed among biotic attacks classes, whatever the organs. Error bars represent standard error. N = 12, 13 and 8 respectively for A, B and C biotic attacks classes.

The quantity of carbon in the reserves was significantly lower in the trees most attacked by insects than in those unaffected and moderately attacked (**FIGURE 6a**). The quantity of reserve carbon in the coarse roots and the stem decreased with the intensity of biotic attacks. The quantity of stored carbon was reduced of 63% for coarse roots and 59% for stem between trees of class A and C. This is particularly clear in the stem, where each biotic attacks class was significantly different from the others. As for NSC concentrations, no difference in the quantity of reserve carbon was observed in the branches among biotic attacks classes.

**FIGURE 6.**
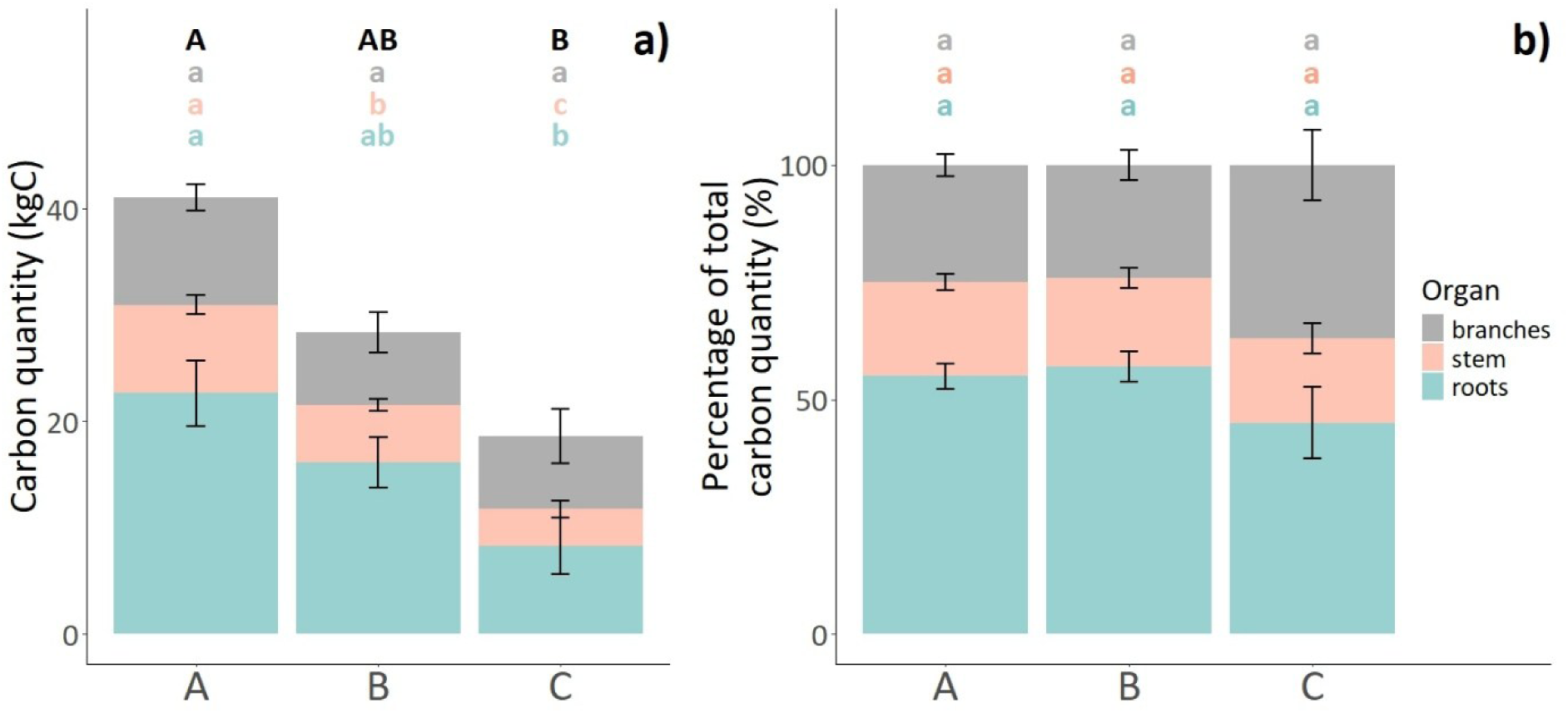
Quantity of carbon reserves at the tree level **a)** in kg by organ (branches, stem, roots), **b)** in proportion of total tree biomass according to the biotic attacks class (A = not significantly attacked; B = moderately attacked; C = heavily attacked). Different lowercase letters represent a significant difference (p < 0.05) among biotic attacks classes for a given organ (the colour of the letter corresponding to that of the organ). Error bars represent standard error. N = 12, 13 and 8 respectively for A, B and C biotic attacks classes.

In the beech trees studied, the coarse roots were the main organ for storing carbon reserves (over 55% for classes A and B and 44% for class C), followed by the branches (25% for classes A and B and 36% for class C) and the trunk (around 20%) (**FIGURE 6b**). The relative importance of these organs for the storage of reserves did not vary significantly among biotic attacks classes.

The quantity of carbon allocated to fruiting tended to increase with the intensity of biotic attacks (11.7, 13.6 and 16 kg of carbon, corresponding to a 37% increase from A to C classes), and to decrease for growth 7.9, 8.4 and 3.8 kg of carbon, corresponding to a 52% decrease from A to C) (**FIGURE *7***). However, these trends were not significant (p = 0.06 between classes B and C for growth).

**FIGURE 7.**
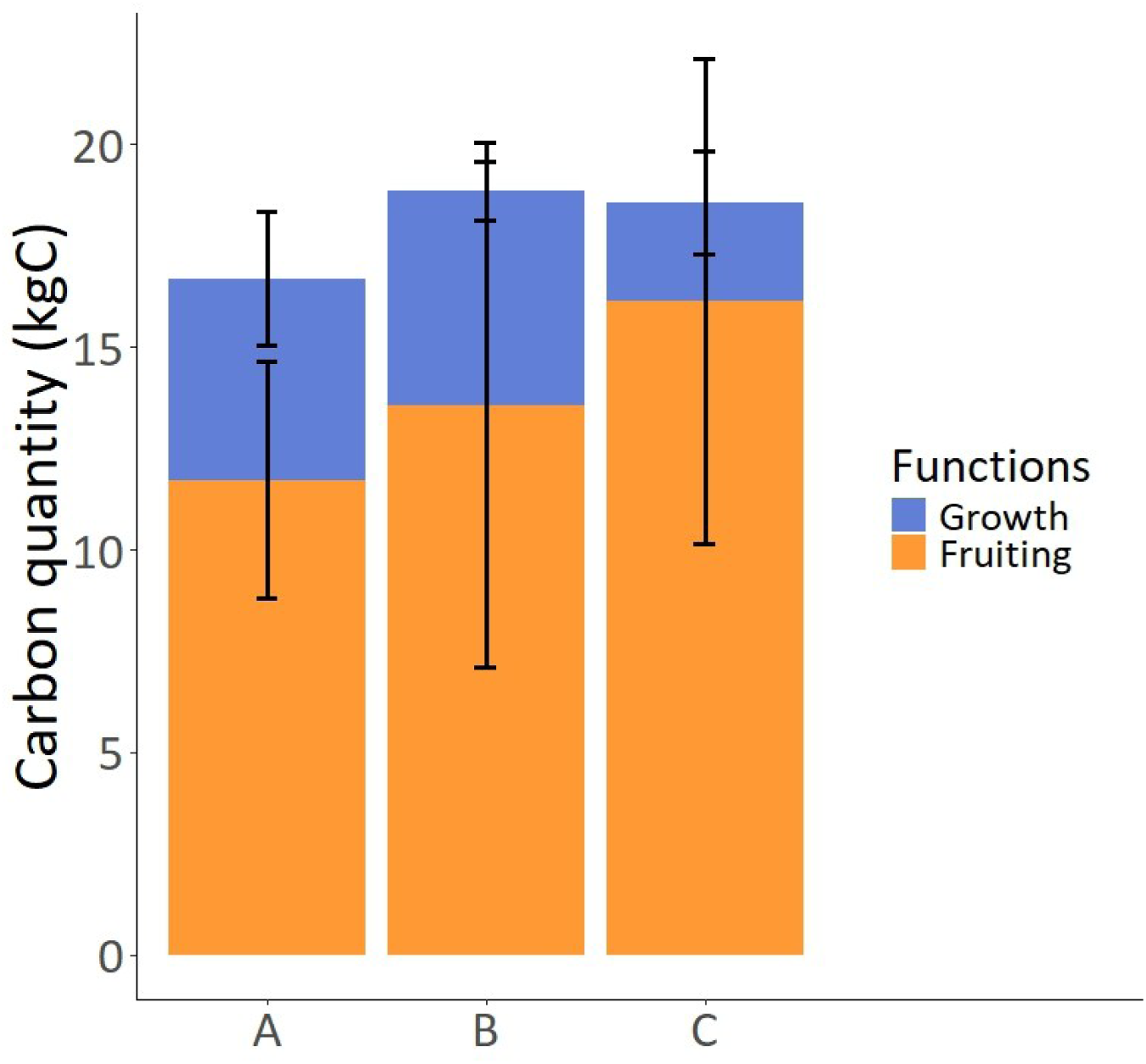
Quantity of carbon at the tree level in kg allocated to aboveground secondary growth and fruiting according to the biotic attacks class (A = not significantly attacked; B = moderately attacked; C = heavily attacked). No significant difference has been observed among biotic attacks classes, whatever the function. Error bars represent standard error. N = 9, 8 and 5 respectively for A, B and C biotic attacks classes.

## 4 Discussion

### 4.1 Carbon invested in secondary stem growth and reserves stock decreased after three consecutive years of extreme soil water deficits

Focusing on non-attacked trees (Class A) allows us to study only the effect of the 2018-2020 drought episode on radial growth and NSC stock. The average growth index in 2020 is 0.36, meaning that radial growth during this year is reduced to about one-third of the average radial growth rate for beech trees in the study site. The same trend is observed for NSC concentrations, it is only 1.5 g.100 g^-1^_DM_ in the first ten centimetres of the trunk at a height of 0.3 m and 3.9 g.100 g^-1^ in the large roots. Such low NSC concentration were reported in severely declining beech trees (Gérard and Bréda 2014). Barbaroux et al. (2003) measured concentrations of 4 and 9 g.100 g^-1^ in the same compartments of healthy beech trees, respectively. By extrapolating the results of Barbaroux et al. (2003) to trees with biomass equivalent to those in our study (the proportion of heartwood in large beech trees is similar or slightly higher than that in smaller trees (Granier et al. 2000; Gebauer et al. 2008), which makes this extrapolation possible), the amount of carbon stored would be approximately 50 kg in the trunk and 32.5 kg in the roots, compared to 6 and 16 kg respectively in the beech trees studied in our study site. Therefore, the radial growth and the amount of carbon reserves measured in our beech trees exposed to three years of intense soil water deficits, are considered to be lower than what would be expected in healthy and non-stressed trees.

Changes in carbon allocation are commonly observed during drought conditions, with metabolism and storage being favoured at the expense of growth (Hartmann and Trumbore 2016; Stefaniak et al. 2024). Particularly in beech trees, radial growth decreases when rainfall is low and temperatures are high from May to July, with a lag-effect the following year (Badeau 1995; Lebourgeois et al. 2005; Scharnweber et al. 2011). The sharp decline in radial growth in 2020 results therefore from both the direct and lagged impact of 2018-2020 soil water deficits. Furthermore, NSC concentrations are generally maintained in the short term following drought (Chuste et al. 2020; Piper and Paula 2020). But during repeated drought episodes, the carbon assimilation due to stomatal regulation is limited. Moreover, the 2018’s soil water deficit induced premature leaf fall in many beech forests in central Europe (Frei et al. 2022), limiting the season for carbon assimilation. Such an event, already described in 2003 in the same region, induced a significant decrease in starch content in beech trees in autumn (Bréda et al. 2006). Thus, the amount of assimilated and stored carbon can be exceeded by the carbon needs of the tree, leading to a situation of carbon starvation after three years of recurrent drought. This would make it impossible to maintain reserves at a normal level (Trugman et al. 2018), and potentially induce tree decline or death (McDowell et al. 2008). Chuste et al. (2020) observed such a decline in juvenile beech trees after three years of extreme water deficits, and a collapse of starch reserves in the trees that died. This decline in NSC concentrations confirms that even non-attacked beech trees show signs of high physiological stress after the 2018-2020 dry episode.

### 4.2 A low tree carbon status: cause or consequence of biotic attacks

In this study, the beech trees with severe biotic symptoms showed both a decrease in radial growth and in their carbon reserves. However, it is difficult to determine whether the insects attacked the more vulnerable trees with limited growth and reserves, or whether the biotic attacks themselves caused a sharper decline in carbon reserves.

The beech trees heavily attacked by insects in 2020 had a lower radial growth than that of undamaged trees since 2009. The cause of the beginning of this decline is not clear and is probably not a consequence of past drought events, the 2003’s drought occurring too many years before the decline in radial growth and no major water deficits occurred in the following years. Thus, excessive rainfall during 2007’s growing season (as reported by Gérard and Bréda (2014) and/or possible defoliation by caterpillars, as observed in the region in 2009 and the subsequent years (Caroulle 2009; Caroulle 2010; Caroulle 2011), could have initiated this growth decline. Whatever the cause of the decline in radial growth, several studies have shown that certain secondary pests prefer to attack trees with low past radial growth (Durand-Gillmann et al. 2014; Stephenson et al. 2019). Indeed, in certain tree species, the level of chemical defence is positively correlated with radial growth (Bazzaz et al. 1987; Mason et al. 2019). In addition, NSC reserves contribute to the production of induced defence compounds (Wiley et al. 2016; Roth et al. 2018). A decrease in carbon reserves could therefore limit the defensive capabilities of trees exposing them to a greater risk of insect attacks. It has been observed that oak trees with lower starch reserves during winter were more susceptible to buprestids attacks the following spring (Dunn et al. 1987; Dunn and Potter 1990). Insects would thus preferentially select vulnerable trees with weakened defences, which would increase the likelihood of successful attacks. This should be the case of *Agrilus viridis* and *Taphrorychus bicolor*, which are considered as secondary pests (Lakatos and Molnár 2009; Brück-Dyckhoff et al. 2019). These findings suggest that NSC reserves in our beech trees may have begun to decline before the bark beetle attacks.

However, It cannot be ruled out that these attacks may still have contributed to the decline in NSC reserves, as observed in several studies (Wiley et al. 2016; Erbilgin et al. 2021). Furthermore, the NSC reserves of the most severely affected trees are so low that it is unlikely that they have reached this level for several years. Indeed, a recent study (Gaertner 2025) has shown that at the level of NSC concentrations measured in the most attacked trees (0.61 g.100 g^-1^) in the base of stem, the short-term survival of beech trees is severely compromised, with a 50% probability of mortality within two years at a concentration below 0.63 g.100 g^-1^. Moreover, the crown condition is similar between trees in classes B and C, suggesting that a deterioration in crown condition alone is not sufficient to cause such a decline in NSC concentrations. Secondary pests of beech trees may also cause a decline in carbon reserves and accelerate tree mortality, directly by feeding on the phloem and the sugars it contains and/or indirectly by inducing the use of carbon reserves for secondary metabolites production for induced defence. However, further studies involving multi-year monitoring of carbon reserves and biotic attacks are needed to confirm this hypothesis. Indeed, beech trees showed active galleries and emergence holes from 2019, and perhaps earlier, so the moment the biotic attacks began and the dynamics of the infestation are unknown. Multi-year monitoring would thus make it possible to determine whether NSC reserves decline as soon as biotic attacks begin or only after a certain level of infestation has been reached.

Furthermore, fruiting and growth are both sustained with new assimilates in European beech (Hoch et al. 2013), and compete for this resource when carbon assimilation is reduced, a steeper growth decline being observed when the drought coincides with a mast year (Hacket-Pain et al. 2017). Thus, high reproductive effort may also limit the carbon quantity allocated to storage, which could have contributed to the decline in carbon reserves, especially in the most attacked trees.

In addition, biotic attacks mainly occurred in the branches and upper part of the stem, but NSC concentrations decreased mainly in the coarse roots and stem. This could suggest a general remobilisation of NSC reserves from long-distance perennial organs towards branches and an impairment of photosynthates redistribution from the leaves. Indeed, phloem transport slow down during drought (Dannoura et al. 2019; Hesse et al. 2019; Salmon et al. 2019) and cambium-feeding insects could impair NSC redistribution via phloem girdling (Paine et al. 1997; Erbilgin et al. 2021). Despite lower concentrations in different organs, our declining trees present similar vertical and radial profile of NSC concentrations to healthy trees from literature (Barbaroux and Bréda 2002; Barbaroux et al. 2003; Hoch et al. 2003). But in our study, coarse roots represent the main organ of storage in carbon quantity instead of stem as usually observed in healthy trees (Barbaroux et al. 2003). Although we were unable to take into account any potential root mortality that would reduce the amount of carbon stored in coarse roots, this compartment seems to constitute the ultimate carbon reserve, the depletion of which would indicate particularly intense carbon stress and a high risk of mortality. Consequently, the measurement of NSC concentrations at the base of the stem (close to the root compartment, but more easily accessible than it) seems to be particularly pertinent to follow carbon status of beech trees.

### 4.3 Carbon allocation to fruiting in declining beech trees

The year 2020 is considered as a mast year for beech in the north-eastern France. Carbon allocation for fruiting in 2020 does not appear to have been affected by the 2018-2020 drought, beechnuts were collected on most of the sampled trees. According to dendrometric inventories, the average density of the study plot is 100 stems per hectare, so the amount of carbon allocated to beechnut and fruits husks production at the stand level is approximately 116 gC.m^-2^. Genet et al. (2010) measured beechnut production ranging from 101 to 129 gC.m^-2^ in mature beech stands during a mast year. In Germany, fruiting accounts for between 100 and 130 gC.m^-2^ in years of abundant beechnut production (Mund et al. 2010). In our study, beech trees did not allocate less carbon to fruiting following the 2018-2020 drought. On the contrary, they may have allocated more carbon to fruiting at the individual level, as stem density was lower in our study site than at the site studied by Genet et al. (2010). Hesse et al. (2021) also observed no decrease in beechnut production in trees under water stress compared to control trees. A recent study showed even a trend to an increased beechnut production with climate warming for forty years (Hacket-Pain et al. 2025). Furthermore, extreme soil water deficits or heat waves can cause beechnut to abort (Nussbaumer et al. 2020). Although nothing caught our attention at the time of harvesting, we did not attempt to distinguish between aborted and healthy beechnuts during harvesting, so it is possible that the allocation of carbon to fruiting is slightly overestimated.

### 4.4 Ecological implications of beech tree’s carbon allocation strategy face to soil water deficit and insect attacks

Since trees cannot escape the constrains to which they are exposed, their survival necessarily depends on their physiological ability to cope with them. This involves adjustments to carbon allocation, which, according to Wiley and Helliker (2012), are a consequence of the evolutionary history of trees. Thus, if carbon starvation during drought is a major threat to tree survival, large carbon reserves and mechanisms favouring the allocation of carbon to sinks necessary for short-term survival would have been selected. Maintaining growth in conditions of water deficit is likely to be detrimental to trees in cases of moderate to severe stress, as it would expose them to greater water stress due to overconsumption of water resources and to the risk of carbon starvation (Wiley and Helliker 2012; Reich 2014), thereby negating the potential benefit in terms of competition (Grime 1977). Maintaining carbon reserves, on the other hand, would be essential for survival, as these reserves fuel respiration (Hartmann and Trumbore 2016), maintain osmotic pressure (Sapes et al. 2021), probably keep the phloem functioning during drought (Sala et al. 2010; Dannoura et al. 2019) and supply tree defence metabolism face to insect attack (Huang et al. 2020). Their decline would therefore only occur in the event of prolonged stress and overconsumption of reserves (Trugman et al. 2018). The depletion in carbon reserves observed in our study suggests that regulatory mechanisms have been overwhelmed by the cumulative constrains such as water deficit and insect attacks. However, whereas carbon reserves deplete, the continued allocation of carbon to fruiting raises questions, as this is not essential for the tree’s short-term survival.

Fruit production can represent a very significant carbon sink (Mund et al. 2010), particularly in mature beech trees during mast seeding years (Genet et al. 2010). In fact, a much more pronounced decline in radial growth is observed during drought when it occurs at the same time as a full mast year (Hacket-Pain et al. 2017). Maintaining a high allocation of carbon to fruiting when assimilation is limited would therefore reduce the amount of carbon available to fulfil reserves or produce defence compounds. Lauder et al. (2019) described these trade-offs in the ‘fight or flight’ conceptual framework. Trees following the ‘fight’ strategy would sacrifice reproduction in favour of functions more useful for short-term survival, such as growth or defence. On the contrary, those following the ‘flight’ strategy would maintain or even increase the allocation of carbon to reproduction at the expense of other sinks, which could compromise tree’s survival. *Fagus sylvatica* thus appear to follow a ‘flight’ strategy, which seems disadvantageous for a long-lived species. Such a strategy could be the result of the reproductive behaviour of beech trees, a masting species. As reproduction can be a rare phenomenon in masting species (Fernández-Martínez and Peñuelas 2021), taking advantage of the most favourable conditions to maximise offspring production is undoubtedly advantageous in terms of fitness. A high priority for carbon allocation to reproduction would then have been selected. In beech trees, the intensity of reproduction is determined by the climatic conditions of the previous two years (Matthews 1955; Piovesan and Adams 2001), beechnut production would remain a priority sink even if it coincides with a dry year as also observed in oak trees, another masting species (Wiley et al. 2017, Bogdziewicz et al. 2020). In the context of climate change, where the frequency and intensity of droughts would increase (Trenberth 2011; Dai 2013), this strategy could be maladaptive. Indeed, we are already seeing a disruption in beech reproduction across Europe, with reproductive effort increasing while reproductive efficiency has declined due to less effective pollination and increased predation (Bogdziewicz et al. 2020; Foest et al. 2024; Hacket-Pain et al. 2025). In addition, beech regeneration succeeds better in semi-shaded conditions (Wagner et al. 2010) because seedlings are sensitive to heat and drought (Muffler et al. 2021). Despite significant investment in reproduction, it is therefore possible that beech forests will experience difficulties to regenerate in the future, especially if sanitary cuts expose the soil and/or if the health of the forests deteriorates, creating a brighter microclimate under the canopy.

Furthermore, our results suggest that secondary pests may contribute to the decline of beech trees by interacting directly or indirectly on carbon allocation. Nowadays, the populations of *Agrilus viridis* and *Taphrorychus bicolor* do not appear to be at epidemic levels, unlike those of *Ips typographus*, which pose a major threat to forest stands. However, we do not know how these populations will react if the frequency and intensity of hot summers and droughts increase in the coming years. Furthermore, we have little knowledge of the impact of silvicultural practices on these insect populations. As *Taphrorychus bicolor* is more abundant in older stands (Špoula et al. 2024), and direct exposure of the bark to light and heat promotes the development of *Agrilus viridis* (Brück-Dyckhoff et al. 2019), it is possible that sudden exposure of mature beech stands to light may be detrimental to trees by exposing thin-barked trunks to heat. Even though these pests are considered as ‘secondary’, it is therefore important to continue studying their biology, population dynamics and impacts on forest ecosystems and trees, as these remain poorly understood.

## 5 Conclusion

In this study, we aimed to investigate the cumulative impacts of the extreme drought episode 2018-2020 and cambium-feeding insects on the carbon allocation in declining beech trees. Regardless of the level of biotic attacks, declining beech trees showed low radial growth and reduced NSC reserves whereas carbon allocation to fruiting was maintained. Beech trees most severely attacked by insects showed a lower carbon status (lower radial growth for ten years and lower carbon reserves) than unattacked trees. However, further investigation is needed to know whether the insects are directly responsible for the decline in carbon reserves, or whether they attacked beech trees with already low carbon reserves. Nevertheless, this study confirms the role of *Agrilus viridis* and *Taphrorychus bicolor* as secondary pests, as these species take advantage of the weakening of trees to colonise them, thereby accelerating their decline. Although we observed signs of physiological decline in the trees most severely affected by insects, we do not know whether this was accompanied also by a decline in their defensive capabilities. The importance of carbon allocation to defence is still poorly understood, particularly in angiosperms, and warrants further investigation.

## Supporting information

All Supplementals

## Acknowledgements

This study was financed by grant from French National Research Agency (ANR) as part of the “Investissements d’Avenir” program (ANR-11-LABX-0002-01, Lab of Excellence ARBRE, Dep-Hetre project). This work was also supported by the Resil Grand-Est project (N°12000767) funded by the French Ministry of Agriculture, Food and Forestry. Pierre-Antoine Gaertner, received a PhD grant from INRAE, supplemented by funding from Labex ARBRE (RESIL Hetre project 24PN09) and the Grand-Est Region (RESIL project, N°12000843).

We thank the French National Service of Forests (ONF) and the municipality of Bliesbruck to allow us to sample beech trees. We thank Erwin Thirion, Patrick Behr and Thierry Paul for their help with fieldwork; and Jade Stucky and Iris Gouchina for their contribution to NSC analyses. We thank the SILVATECH platform, ISC from UMR 1434 SILVA, 1136 IAM, 1138 BEF, 4370 EA LERMAB Research Center INRAE Nancy-Lorraine for its contribution to the NSC analyses and for welcoming and training students. ChatGPT was also used to translate and improve the syntax of some sentences.

## Author contributions

NB and CM contributed to the experiment design. JL, HS, TL, VB and CM took part in field sampling. HS and TL conducted the crown transparency assessment and the record of insect attacks. BG conducted the NSC analysis. JL, PAG and CM performed soil sampling, JL conducted soil data analysis and BILJOU© simulations. PAG conducted tree core preparation and tree ring measurements. PAG performed the data analysis and wrote the first draft of the manuscript. PAG, NB, BG, JL, VB, FXS and CM contributed to editing and revising the manuscript.

## Data availability

Authors agree to make data available on a public repository.

