## Supplementary material for "Impact of secondary pests on carbon allocation in declining beech trees after the 2018-2020 drought episode": All Supplementals

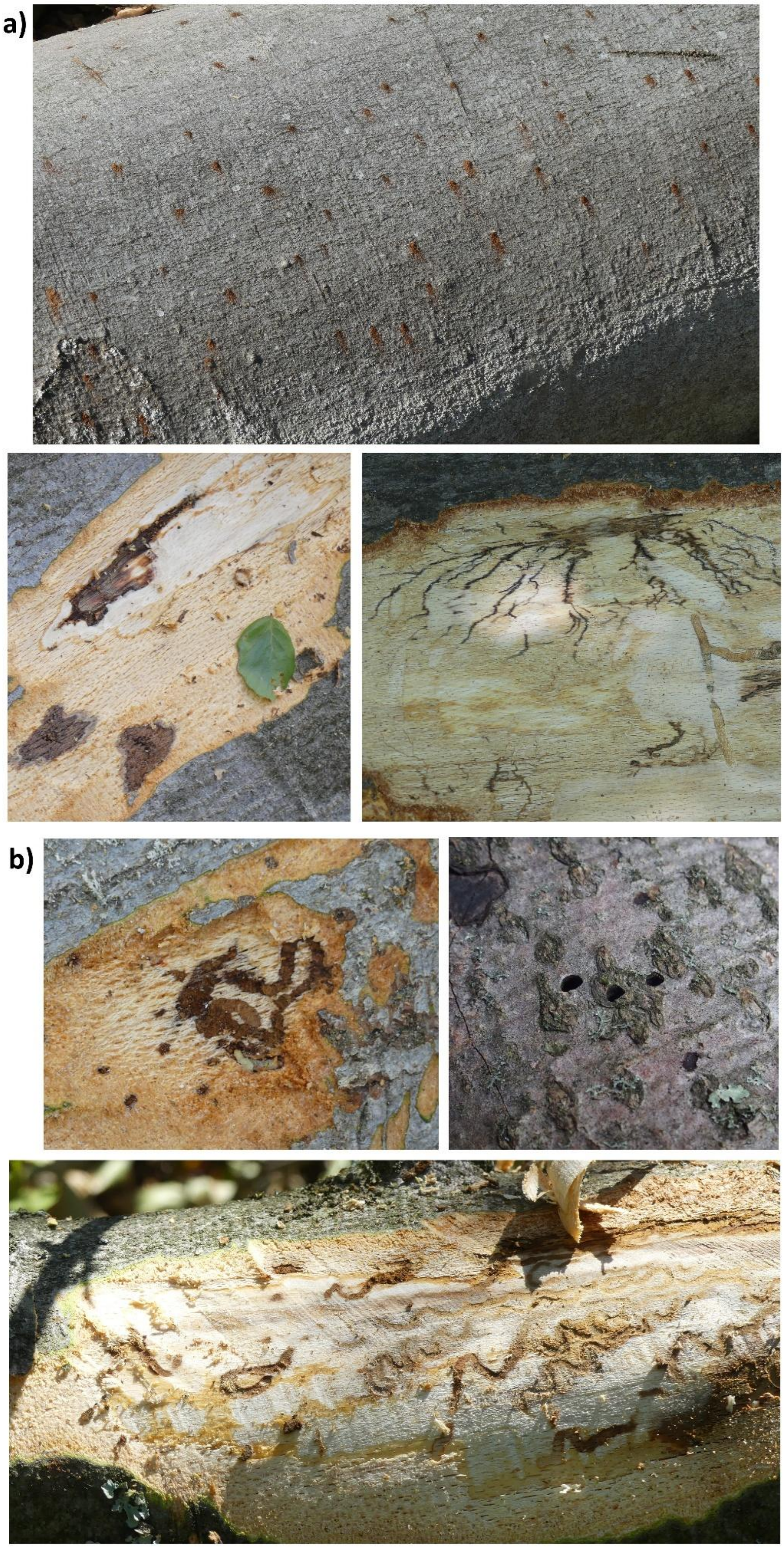


**FIGURE S1** Galleries and emergence holes of **a)** *Taphrorychus bicolor* and **b)** *Agrilus viridis* ©Hubert Schmuck.

**S2: Estimation of the quantity of carbon stored and of the quantity of carbon allocated to aboveground radial growth and reproduction**

### 1. Biomass estimation for each compartment

The first step before calculating the quantity of carbon allocated to different function was to estimate the biomass of the sampled organs.

The biomass of coarse roots (1) and branches with a diameter below 4cm (2) were estimated using allometric equations based on tree diameter and height for European beech from Genet et al. (2011). For branches, the biomass calculated using the allometric equation was weighted with branch mortality to avoid overestimating living branch biomass of declining trees (the biomass calculated using the allometric equation was multiplied by 105 – percentage of branch mortality (BM) (BM was subtracted to 105 instead of 100 because this note was assessed in 10% non-centred steps, which means that a BM note of 100% represents a branch mortality between 91 and 100%, so it doesn't necessarily mean that there is no living branches)).

The biomass of twigs was estimated using the equation from Ottorini & Le Goff (1998) in order to calculate the proportion of twigs in the total biomass of branches with a diameter below 4cm (equation (1)). This proportion was estimated at 1.6% and was used in the calculation of the quantity of carbon from NSC concentrations in branches and twigs.

$$(1)\ \ biomass_{coarse\ roots} = 0.08 + 50.4 \cdot (d^2 \cdot h)^{1.13} + 16.3 \cdot (h \cdot d^2) \cdot 1.31$$

$$(2)\ \ biomass_{branches} = (38.5 \cdot (d^2 \cdot h)^{0.8} + 21.8 \cdot (h \cdot d^2) \cdot 0.8) \cdot (\frac{105-BM}{100})$$

$$(3)\ \ biomasse_{twigs} = -7.8248 + 2.0820 \log{(C_{130})}$$

with d, the diameter at breast height; $C_{130}$, the circumference at breast height; h, the tree height; and BM, the percentage of branch mortality.

Stem biomass was estimated as the product of stem volume and wood density. According to Barbaroux et al., (2003), beech wood density is very homogenous from a ring to another one. So, wood density was set to 566 g dm$^{-3}$. Stem was considered as a truncated cone divided in two sections (Barbaroux et al., 2003): the first section from 1.3m to 11m, and the second section from 11m to 22m. The biomass for each section was calculated using equation (4):

$$(4)\ \ biomass = \frac{\pi}{12} \cdot l \cdot ({d_b}^2 + d_b \cdot d_t + {d_t}^2) \cdot 566$$

with $d_b$, the diameter at the base of the stem section; $d_t$, the diameter at the tip of the stem section; l, the length of the stem section; and 566g dm$^{-3}$, the density of beech wood.

The collected branches and their beech nuts and vegetative infrutescence tissues were weighed fresh, and a sample of these parts was taken and weighed. These samples were dried in an oven at 80°C for three days, then weighed to determine their moisture content and to establish an allometric relationship between the dry weight of the branches and the total dry weight of beech fruits. The method for estimating the weight of beech fruits production per tree is derived from (Barbaroux, 2002). A relationship between fresh weight and dry weight of the branches samples was first set ($R^2$ = 0.97):

$$(5)\quad dry\ weight = 0.727 \cdot fresh\ weight - 2.493$$

This made it possible to calculate the total dry weight of the sampled branches and then to calculate the ratio of the dry weight of the harvested beech fruits and fruits husks to the dry weight of the branch from which they were taken; this was then multiplied by the total biomass of the tree's living branches with a diameter below 4cm to obtain the dry of beech nuts and husks per tree. Flowers and catkings disappeared at harvest time and were not quantified. This flowing part of reproduction represents less than 1% of annual production (Lebret et al., 2001).

2. <u>Carbon quantity stored</u>

Once the biomass of each organ has been calculated, the method for estimating the amount of carbon stored per tree (in gC) involves multiplying the biomass of each organ in the tree by its NSC concentration, expressed as glucose equivalent, and by 0.4, which corresponds to the proportion of carbon in glucose.

<u>Roots:</u> As the coarse roots have not been divided into sub-compartments, the method was applied as it stood.

<u>Branches:</u> As the branches exhibits significant differences in NSC concentrations between the base and the twigs from the last growing season (Barbaroux et al., 2003), the NSC concentrations were weighted by 1.6%, which is the estimated proportion of twigs in the total biomass of branches over 4 cm in diameter. The NSC concentrations in the twigs were calculated by averaging the concentration in the twigs from the growing seasons 2019 and 2020. We also assumed that the NSC concentrations vary little in branches between a 2.4cm-diameter section and a 4cm-diameter section, enabling us to use the corresponding equation in Genet et al., (2011). Finally, the NSC concentrations of the branches were calculated by taking the weighted average of the NSC concentrations in branches and twigs

(coefficients of 98.4 and 1.6 respectively), then multiplying this by the total biomass of the branches and 0.4 to convert it to carbon content (in gC).

Stem: The stem was divided into two sections (1.3–11 m and 11–22 m), each of which was further divided into four sub-compartments corresponding to the radial sections of 2-cm-thick core samples (0–2 cm, 2–4 cm, 4–6 cm, 6 cm–core). For the base, core sections deeper than 6 cm were combined and their concentration calculated by averaging the NSC concentrations weighted by the cross-sectional area of each section. As the NSC concentrations were considered to vary linearly along the trunk section, the NSC concentrations at the base and top of each trunk section were averaged to calculate the concentration of each sub-compartment. The biomass of each sub-compartment was calculated using equation (6).

$$(6)\quad biomass_{section} = \pi \cdot l \cdot (d_b - d_t - 2 \cdot (2 + n)) \cdot 566$$

with $d_b$, the diameter at the base of the stem section; $d_t$, the diameter at the tip of the stem section; l, the length of the stem section; n, the number of the core section (from 1 for the 0-2cm section to 4 for the 6cm-core section); and 566g $dm^{-3}$, the density of beech wood.

For each sub-compartment, the biomass was then multiplied by the NSC concentrations in glucose equivalents and by 0.4 to convert it to carbon content (gC). Finally, the carbon content of all sub-compartments in the two sections of the stem was summed to calculate the carbon content stored in the stem.

#### 3. Carbon quantity in fruits

To estimate the amount of carbon allocated to fruiting, the dry weight of beech nuts per tree was multiplied by the proportion of carbon in beech nuts (50%, (Lebret et al., 2001)).

#### 4. Carbon quantity in annual radial growth

To estimate the amount of carbon allocated to aboveground radial growth during a year, the same method used to estimate stem biomass was employed. The principle for calculating stem volume increment in 2020 ($V_{2020}$) is to subtract the volume of the stem without the 2020 growth ring from the total stem volume. Although Pressler's law is not strictly verified for beech trees sampled at 1.3 and 11 m height, no significant difference in BAI was observed between the cores sampled at 1.3 m and 11 m height in 2020 (Fig. S2.1). Pressler's law therefore appears to be a sufficiently accurate approximation of reality for the year 2020, and the ring area was considered constant from 1.3 m to 22 m in height. The volume increment in 2020 ($V_{2020}$) was thus calculated using equation (7) by adapting equation (4).

$$(7)\quad V_{2020} = \frac{\pi}{12} \cdot l_{stem} \cdot \left(d_{1.3m} \cdot d_{22m} - 8 \cdot \frac{BAI_{2020}}{\pi} - 4 \cdot \sqrt{\left(\frac{d_{1.3m}}{4} - \frac{BAI_{2020}}{\pi}\right) \cdot \left(\frac{d_{22m}}{4} - \frac{BAI_{2020}}{\pi}\right)}\right)$$

with $V_{2020}$, the stem volume increment in 2020; $d_{1.3m}$, the diameter at breast height; $d_{22m}$, the diameter at 22m height; $l_{stem}$, the length of the stem; and $BAI_{2020}$, the basal area increment in 2020.

This volume was then multiplied by the density of the wood (566 g $dm^{-3}$) and the proportion of carbon in the biomass of beech stem (46%) (both values are taken from (Barbaroux et al., 2003)) to calculate the amount of carbon allocated to aboveground radial growth.

**S3: Standardisation of the radial growth according to tree age and to the height of sampling**

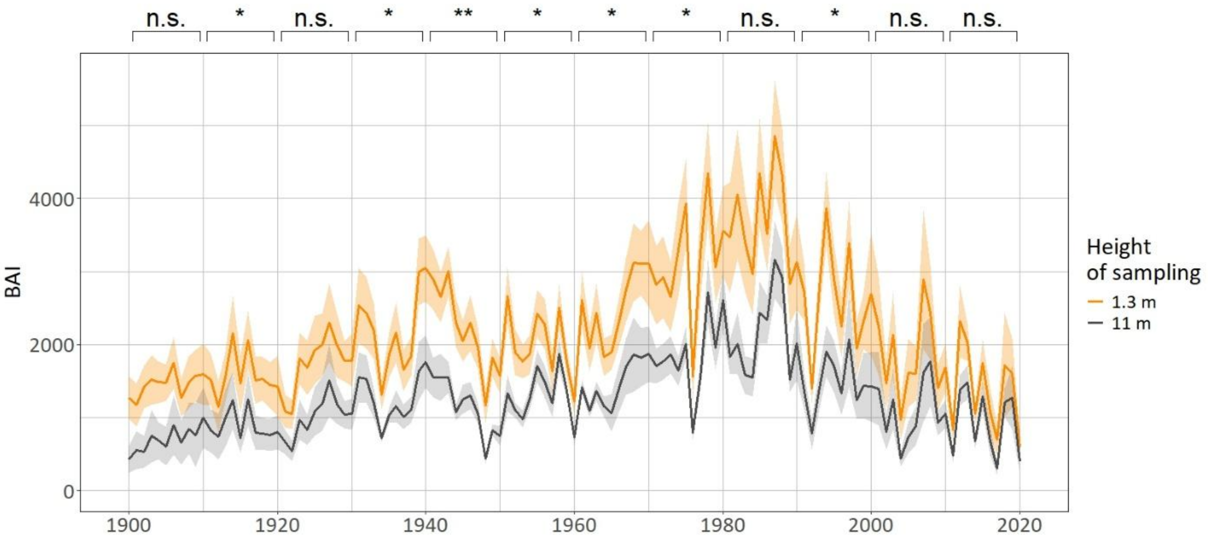


**FIGURE S3.1** Basal area increment (BAI, in mm²) from 1900 to 2020 as a function of the height of sampling for the eight trees sampled at both 1.3 and 11 metres. Brackets represent significant differences in averaged BAI (BAI was averaged for each decade and for each tree) among heights of sampling at 1.3m and 11m. (p > 0.05 : n.s. ; p < 0.05 : * ; p < 0.01 : **).

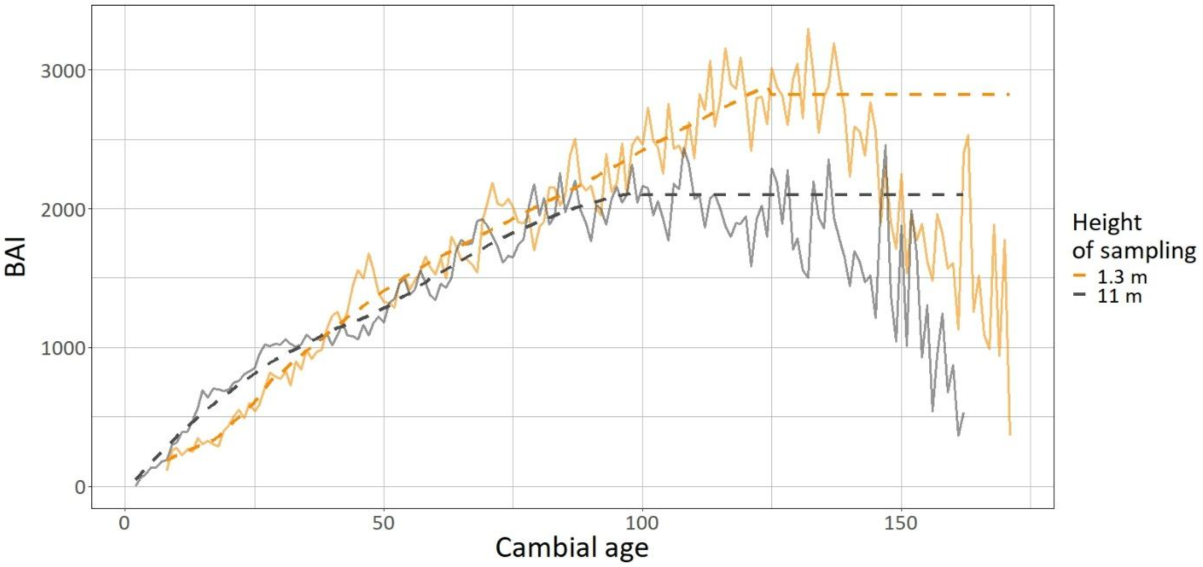


**FIGURE S3.2** Basal area increment (BAI, in mm²) as a function of cambial age for each height of sampling. All the trees sampled for the dendrochronological analysis were used to produce this graph. Continuous lines represent the mean basal area increment for each height of sampling, dashed lines represent the LOESS fit for the BAI ~ cambial age relationships for each height of sampling.

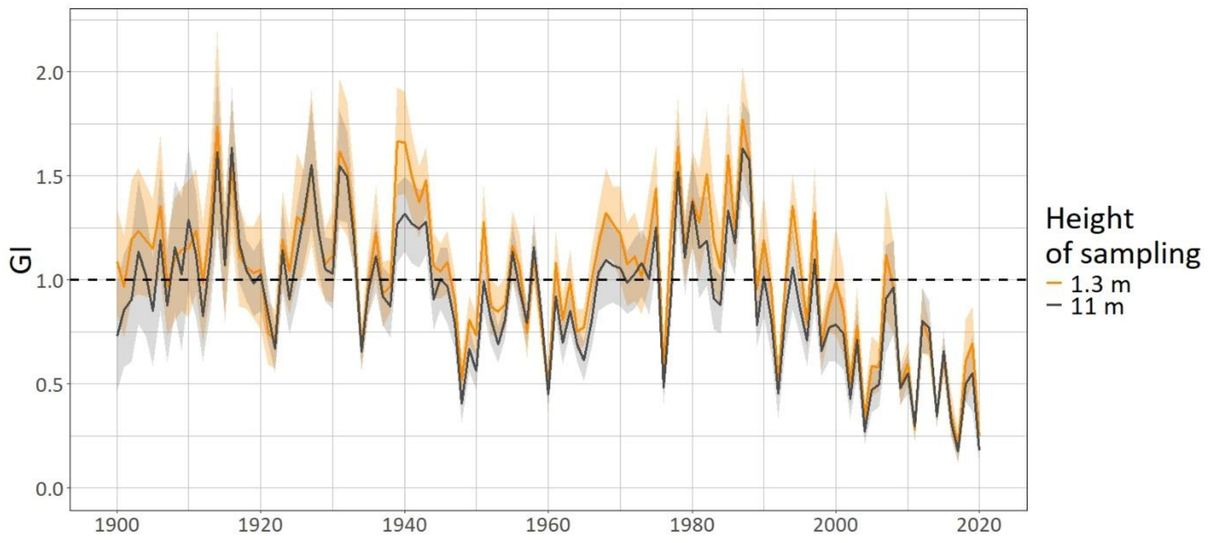


**FIGURE S3.3** Detrended basal area increment (called Growth Index, GI) from 1900 to 2020 as a function of the height of sampling for the eight trees sampled at both 1.3 and 11 metres.

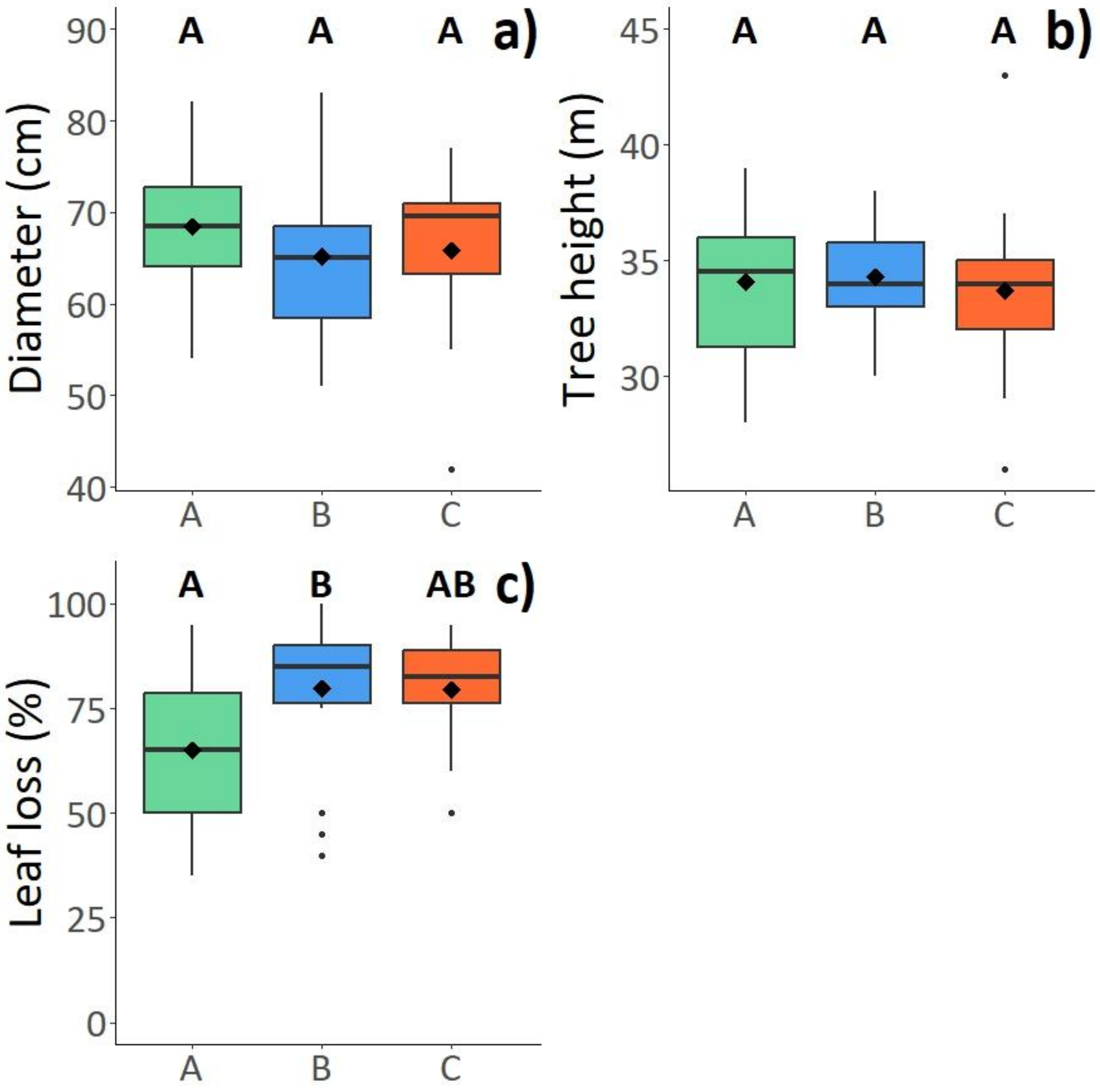


**FIGURE S4 a)** Diameter, **b)** tree height and **c)** leaf loss of the trees sampled in 2020 according to the biotic attack class (A = not significantly attacked; B = moderately attacked; C = heavily attacked). In boxes, diamonds represent the mean, lines represent the median. Different letters represent a significant difference ($p < 0.05$) among biotic attacks classes. N = 18, 18 and 10 from A to C.
